# Identification of genome-wide nucleotide sites associated with mammalian virulence in influenza A viruses

**DOI:** 10.1101/416586

**Authors:** Yousong Peng, Wenfei Zhu, Zhaomin Feng, Zhaozhong Zhu, Zheng Zhang, Yongkun Chen, Suli Liu, Aiping Wu, Dayan Wang, Yuelong Shu, Taijiao Jiang

## Abstract

**Motivation:** The virulence of influenza viruses is a complex multigenic trait. Previous studies about the virulence determinants of influenza viruses mainly focused on amino acid sites, ignoring the influence of nucleotide mutations.

**Results:** We collected more than 200 viral strains from 21 subtypes of influenza A viruses with virulence in mammals and obtained over 100 mammalian virulence-related nucleotide sites across the genome by computational analysis. Interestingly, 50 of these nucleotide sites only experienced synonymous mutations. Further experiments showed that synonymous mutations in the top two of these nucleotide sites, i.e., PB1-2031 and PB1-633, enhanced the pathogenicity of the viruses in mice. Finally, machine-learning models with accepted accuracy for predicting mammalian virulence of influenza A viruses were built. Overall, this study highlighted the importance of nucleotide mutations, especially synonymous mutations in viral virulence, and provided rapid methods for evaluating the virulence of influenza A viruses. It could be helpful for early warning of newly emerging influenza A viruses.

## 1 Introduction

Influenza viruses are a kind of negative-sense single-stranded RNA viruses. There are four types of influenza A viruses: A, B, C and D (Nakatsu, et al., 2018). Among them, influenza A viruses (IAV) can infect a wide range of avian species and some mammal species including humans (Webster, et al., 1992), and cause the greatest morbidity and mortality in human populations (Thompson, et al., 2003). Based on the two surface glycoproteins—hemagglutinin (HA) and neuraminidase (NA), IAV can be divided into different subtypes (Taubenberger and Kash, 2010), such as H3N2, H5N1, and so on. In recent years, more and more subtypes of IAV were reported to cross species and caused human infections (Gao, et al., 2013; Su, et al., 2015; Taubenberger and Kash, 2010), such as H5N1, H7N9, H5N6, H10N8 and so on. How to prevent and control the newly emerging IAV is still a huge challenge for human beings.

Virulence is an important phenotype of influenza viruses. Understanding the mechanism of viral virulence in host, especially in mammals, could help for accurate evaluation on the risk of the virus. Many previous studies showed that the mammalian virulence was attributed to the effects of multiple genes (Chen, et al., 2007; Govorkova, et al., 2006; Lycett, et al., 2009). For example, Lycett et al. identified lots of mammalian virulence determinants (amino acid sites) in multiple proteins of the highly pathogenic avian influenza (HPAI) H5N1 virus by computational analysis of virulence data obtained from more than 20 studies (Lycett, et al., 2009). It was reported that many amino acid sites can influence the viral virulence in mammals (Chin, et al., 2014; Ilyushina, et al., 2010; Kamal, et al., 2014; Lycett, et al., 2009; Otte, et al., 2015; Wang, et al., 2016). Among them, PB2-627 is the most widely recognized virulence determinant (Chin, et al., 2014) and E627K mutation can increase the pathogenicity of influenza viruses in mammalian cells. However, so far, most of the identified virulence determinants were only analyzed from amino acid levels.

Mutations at the nucleotide sites were reported to affect the transcription, translation and packaging process of IAV (Ferhadian, et al., 2018; Kobayashi, et al., 2016). Taking the packaging process as an example, many nucleotide sites in both coding and non-coding regions were found to take part in the packaging process (Gog, et al., 2007; Marsh, et al., 2008; Williams, et al., 2018). In Williams’s study, it showed that synonymous mutations in low-nucleoprotein-binding regions of the genome could impact genome packaging and result in virus attenuation by altering RNA structures (Williams, et al., 2018). Wang et al. and Zhao et al. reported that mutations in the non-coding regions could influence virion incorporation, viral RNA synthesis and protein expression at transcription and translation levels (Wang, et al., 2015; Zhao, et al., 2014). In addition, nucleotide-level mutations can also interfere with antiviral response of host cells by lowering or even abolishing the binding of host microRNAs to the viral mRNAs (Ingle, et al., 2015; Wang, et al., 2017). Based on above results, it is indicated that the mutations in nucleotide sites are extremely likely to affect the virulence of influenza viruses. However, few studies have definitely investigated the influence of nucleotide mutations on viral virulence, let alone the systematic identification of virulence determinants from nucleotide level.

In this study, we collected more than 200 viral strains with virulence in mammals from 21 subtypes of IAV which is the largest dataset ever recorded to our knowledge. Then, we systematically identified genome-wide mammalian virulence-related nucleotide sites by computational analysis. We further validated the top two of these virulence determinants by *in vivo* experiments. Finally, we built computational models with acceptable accuracy for rapidly predicting the virulence of the viruses.

## 2 Materials and Methods

### 2.1 Viral strains with mammalian virulence and with the genomic sequences available from public databases

To obtain the viral strains of IAV with experimentally determined mammalian virulence, we firstly obtained 487 abstracts from PubMed database (Wheeler, et al., 2002) by searching “influenza [TIAB] AND virulence [TIAB] AND ((ferret [TIAB]) OR (mice [TIAB]))” on April 18th, 2017. Then, each abstract was manually screened based on whether it contained viral isolates with mammalian virulence and 157 abstracts were retained. Finally, the full texts of these abstracts were read carefully and 279 viral strains with experimentally determined mammalian virulence were compiled from these papers. Among them, only 228 strains from 92 individual studies had the genomic sequences which can be obtained from the database of Global Initiative on Sharing All Influenza Data (GISAID) (Bogner, et al., 2006) and Influenza Virus Resource (Bao, et al., 2008). Therefore, they were used for further analysis. The mammalian virulence and the accession numbers for genomic sequences of these strains were shown in Table S1 and Table S2, respectively.

According to Lycett’s method (Lycett, et al., 2009), each viral strain was manually classified either as low-virulent or high-virulent in mammals based on the amalgamated experimental evidence. Although the virulence of viral strains were obtained in different animals and using different protocols, the results were generally concordant: 213 strains could be classified clearly either as high-virulent in mice or ferrets, or only caused mild symptoms or few death; 16 strains had virulence data in both mice and ferrets, 13 (81%) of which were identical (with same virulence level in both mice and ferrets).

### 2.2 Data preprocessing

Because of the difficulties in aligning protein or genomic sequences of multiple subtypes of HA or NA, for the surface proteins HA and NA, we only considered H5 and N1 subtypes. The genomic sequences for each segment of viral strains were aligned by MAFFT (version: 7.127b) (Katoh and Standley, 2013) and then checked manually. The coding sequences for ten internal proteins (PB2, PB1, PB1-F2, PA, PA-X, NP, M1, M2, NS1 and NS2) and two surface proteins (H5 and N1) were extracted from the genomic sequence, and then translated into protein sequences with a perl script. All the conserved sites (with only one kind of amino acid or nucleotide) were removed from the protein or nucleotide sequences.

To identify the nucleotide (or amino acid) sites which were associated with mammalian virulence of IAV, the genomic (or protein) sequences of viral strains were converted into a binary matrix according to the method in Lycett’s study (Lycett, et al., 2009). Each row of the matrix represented a viral isolate genome (or proteome), and columns represented nucleotide (or amino acid) sites. An additional column for the virulence phenotype was added with 1 and 0 encoding the high-virulent and low-virulent phenotype, respectively. The binary coding at each variable site was determined with the procedure as in Lycett’s study via a perl script (see Supporting Information for details).

### 2.3 Association of amino acid sites with virulence

The degree of association between the mammalian virulence of viral strains and the amino acid sites in the ten internal proteins (PB2, PB1, PB1-F2, PA, PA-X, NP, MP1, MP2, NS1 and NS2) was analyzed by Fisher’s exact test based on the binary coding of virulence phenotype and amino acid residues, which resulted in a p-value for each site. To correct for multiple comparisons, q-value method was used to control the false positive rate with the help of *qvalue()* function in the R package “qvalue” (Bass, et al., 2015). The amino acid sites with p-value < 0.05 and q-value < 0.05 were considered to be significantly associated with mammalian virulence.

To reduce the influence of sampling bias of viral strains, bootstrap method was used as follows: firstly, a novel data set with the same size as the original data set (228 viral strains) was generated by sampling from the original data set with replacement; then, the association analysis was performed using the above method for the novel data set to identify the amino acid sites which had significant association with virulence (p-value < 0.05 and q-value < 0.05). This process was repeated 1000 times. The number of significant associations between an amino acid site and virulence in the bootstrapping process was defined as the bootstrap value for the site. The amino acid sites identified to be associated with mammalian virulence in the original data were further filtered by the bootstrap value of the site. Only the sites with the bootstrap values greater than 300 were used for further analysis.

Similar methods were used to identify the amino acid sites which were significantly associated with mammalian virulence in H5 and N1 protein of H5NX strains (including H5N1, H5N2,H5N5, H5N6, H5N8 and H5N9) and HXN1 strains (H1N1,H5N1, H6N1 and H7N1). The p-value, q-value, and bootstrap value for all variable amino acid sites in internal and surface proteins were presented in Table S3.

### 2.4 Association of nucleotide sites with virulence

The methods for identifying virulence-related nucleotide sites were similar to those for amino acid sites, i.e., Fisher’s exact test, q-value and bootstrap method. Besides, two additional steps were carried out to eliminate the influence of amino acid sites and codon usage bias. The first step was to remove the nucleotide sites which encoded the mammalian virulence-related amino acid sites. The second step was to remove the codon usage bias using a permutation method as follows: firstly, the frequency distribution of codons encoding each amino acid was derived based on all genomic sequences used in this study; secondly, each genomic sequence was re-generated by codon according to the frequency distribution of codons mentioned above while keeping the protein sequence unchanged; thirdly, the association analysis was conducted as above on the permutated data set to identify the nucleotide sites which had significant association with virulence (p-value < 0.05 and q-value < 0.05). This permutation process was repeated 1000 times. For each nucleotide site, a permutation p-value was calculated by dividing the number of association of the site with virulence by 1000. Finally, only those nucleotide sites with p-value (Fisher’s exact test) < 0.05, q-value < 0.05, bootstrap value > 400 and permutation p-value < 0.05 were selected as mammalian virulence determinants. Details for all variable nucleotide sites in both internal and surface protein-coding genes were listed in Table S4.

### 2.5 Plasmids, mutagenesis, and virus generation

Eight expression plasmids of the 2009 pandemic H1N1 virus A/California/04/2009 (CA, H1N1) or human isolated Eurasian avian like swine influenza virus A/Hunan/42443/2015 (HuN, H1N1) based on the reverse genetic system were constructed by Chinese National Influenza Center. Full-length viral cDNAs were cloned into the plasmid vector pHW2000 as previously reported (Hoffmann, et al., 2000; Hoffmann, et al., 2001). Mutations were introduced into the PHW2000-CA-PB1 or PHW2000-HuN-PB1 plasmid to generate the mutant segment, and were termed as m2031 (A > G on PB1-2031) or m633 (A > G on PB1-633). The presence of the introduced mutations and the absence of additional unwanted mutations were verified by sequencing.

To rescue viruses, eight corresponding reverse-genetics plasmids containing the dsDNA representing each gene segment were co-transfected into monolayers of co-cultured 293T/MDCK cells using the PolyFect (Qiagen) reagent. After 24 hours post transfection, TPCK-Trypsin (0.25μg/ml) were added to support the virus replication. Recombinant wild type viruses (CA and HuN), and their mutants (m2031 and m633) were generated and confirmed by both hemagglutinin test and sequencing.

### 2.6 Studies in mice

All mice experiment protocols were approved by the Ethics Committee of the National Institute for Viral Disease Control and Prevention, Chinese Center for Disease Control and Prevention (20160226008). Groups of 8 to 10-week-old female C57BL/6J mice (five in each group) were anesthetized with 0.2 ml pentobarbital sodium and inoculated intranasally with10^6^ TCID_50_ of the recombinant viruses (with the volume of 50 μl). The control group was inoculated with PBS. Body weight was measured daily for 14 days. Mice that lost more than 30% of their original weight were euthanized for humane reasons.

### 2.7 Predicting mammalian virulence of IAV based on nucleotide and amino acid sites

Because of the difficulties in aligning the protein or genomic sequences of multiple subtypes of HA or NA, we only used the nucleotide and amino acid sites in internal segments to predict the mammalian virulence of IAV. Each nucleotide or amino acid site was considered as an explanatory variable, while the virulence was considered as the response variable. Both of them were binary variables as mentioned above. We firstly assessed the ability of single nucleotide or amino acid site in predicting the virulence. Then, combinations of multiple nucleotide sites and/or amino acid sites were used in the prediction. To make the model as concise as possible, feature selection was conducted by using Weka (version 3.6.1) (Hall, et al., 2009) with the attribute evaluator of “CfsSubsetEval”, the searching method of “BestFirst (options: -D 1 -N 5)” and the attribute selection mode of ten-fold cross-validations. The features appeared in seven or more folds were finally selected. The methods of decision tree (J48), random forest, logistic regression, neural network (MultilayerPerceptron) and naive bayes were used respectively to predict the virulence based on the selected features, i.e., nucleotide sites and/or amino acid sites, with default parameters in Weka. Ten-fold cross-validations were used for evaluating the performance of these models. The Area Under a receiver operating characteristics Curve (AUC), predictive accuracy, sensitivity and specificity were used to measure the performance of models.

### 2.8 Statistical analyses

All the statistical analysis was carried out in R (version 3.2.5) (R Core Team, 2016). Fisher’s exact test was performed with the help of function *fisher.test()*.

## 3 Results

### 3.1 Viral strains with experimentally determined mammalian virulence

A total of 228 viral strains of IAV with genomic sequences that can be gained from public databases and with experimentally determined virulence in mammals (either mice or ferrets) were obtained from 92 studies by manual curation (Table S1&S2). Each viral strain was manually classified either as low-virulent or high-virulent in mammals based on the amalgamated experimental evidence according to Lycett’s method (Lycett, et al., 2009). Specifically, a viral strain was classified as high-virulent if a high mortality was observed with a low dose in infection studies, or the viral strain had a 50% lethal dose of <=10^3^ 50% egg infectious doses (EID) or <= 10^2^ plaque-forming unit (PFU); otherwise, it was classified as low-virulent. 65 viral strains were classified as high-virulent in mammals, and the others were low-virulent. They belonged to 21 subtypes (Table 1) and contained most subtypes of IAV which have been reported to infect humans (marked with asterisks), such as H5N1, H5N6, H7N9, H9N2, and so on. The viral strains of H5N1 subtype, which have caught the attention of the world in the last 20 years (Peng, et al., 2017), accounted for more than 50% of all strains and more than 80% of all high-virulent strains. The virus strains of H9N2 subtype have also been circulating in the last 30 years (Sun, et al., 2010). Although there were more than 20 strains of the subtype, few of them were high-virulent in mammals. Similar situations were observed in most subtypes, such as H7N9, H1N1 and H5N2. In contrast, the H5N8 subtype, which causes widespread outbreaks in the globe in recent years (Lee, et al., 2017), had seven strains and three of them were high-virulent.

**Table 1.**
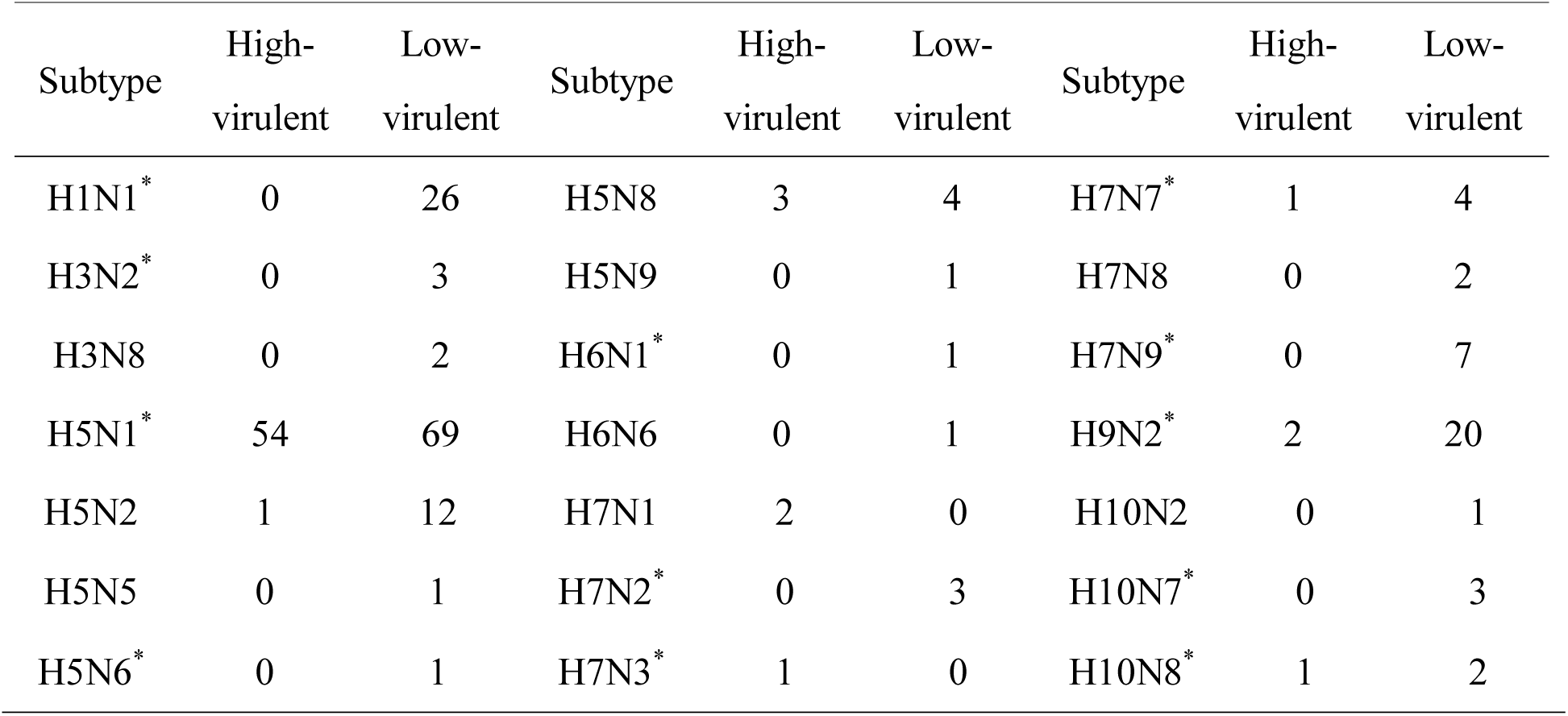
Subtype distribution of the viral strains used in this study. The asterisks refer to the subtypes which have been reported to infect humans.

By isolation host, the viral strains included 131 from the avian, 78 from humans, 13 from swine, 5 from the environment and 1 from tiger. Among them, 25% (33) of avian isolates, 38% (30) of human isolates, 8% (1) of swine isolates and 100% (1) of tiger isolate were classified as high-virulent in mammals.

Then, we investigated the isolation time and place of viral strains. Most of viral strains were isolated from 1996 to 2016, especially in the period from 2003 to 2010 when more than ten strains were isolated each year (Fig. S1). Some isolation peaks could be observed, such as in 1997 and 2013. Most of viral strains were isolated from countries in East and South East Asia (Fig. S2), such as China, Vietnam and Indonesia. The remaining viral strains were isolated from Europe, North America, other regions of Asia, and Australia.

### 3.2 Identification of genome-wide nucleotide sites associated with mammalian virulence

Based on the data collected, we attempted to identify genome-wide nucleotide sites which may be associated with mammalian virulence of multiple subtypes of IAV. For the genes encoding internal proteins, including PB2, PB1, PA, NP, MP and NS, the data of all subtypes of IAV were used for analysis; for the genes (H5 and N1) encoding surface proteins, due to the difficulties of aligning genomic sequences of multiple subtypes of HA or NA, the data of H5NX subtypes, including H5N1, H5N2, H5N5, H5N6, H5N8 and H5N9, and HXN1 subtypes, including H1N1, H5N1, H6N1 and H7N1, were used. A computational work flow shown in Fig. 1 was developed to identify mammalian virulence determinants at the nucleotide level.

**Fig 1.**
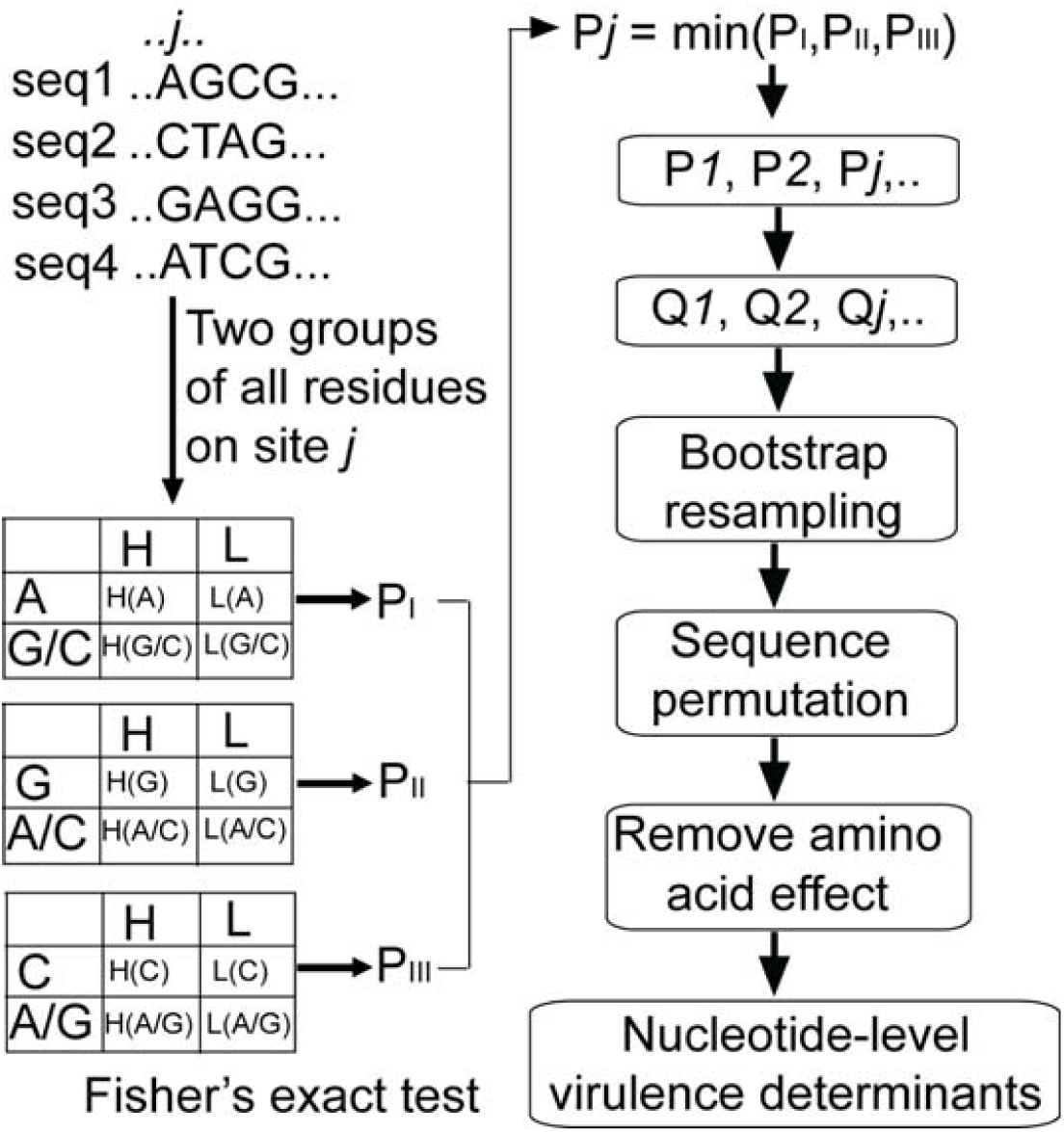
The work flow of identifying nucleotide sites associated with mammalian virulence. Please see Methods for details.

Firstly, according to Lycett’s method (Lycett, et al., 2009), Fisher’s exact test was used to analyze the association between the nucleotide sites in the genome of IAV, and the mammalian virulence (Fig. 1). Then, q-value method was used to control the false positive rate. To reduce the influence of sampling bias of viral strains and to remove the codon usage bias, bootstrap method and permutation were respectively conducted (see Methods). And finally, the effect of mammalian virulence-related amino acid sites should be removed. This was achieved by firstly identifying the amino acid-level mammalian virulence determinant with the methods of Fisher’s exact test, q-value and bootstrap as mentioned above (see Methods). Fifty-eight amino acid sites were found to have significant associations with mammalian virulence of IAV (Table S3), including 40 sites in ten internal proteins (PB2, PB1, PB1-F2, PA, PA-X, NP, MP1, MP2, NS1 and NS2) and 18 sites in surface proteins (H5 and N1). Some of these amino acid sites were reported to influence the pathogenicity of IAV in mammals in previous studies, such as PB2-627 (Chin, et al., 2014), NS1-195 (Bornholdt and Prasad, 2008), H5-279 (Wu, et al., 2008), and so on. The nucleotide sites encoding these 58 amino acid sites were removed for the further analysis.

Based on the work flow in Fig. 1, 137 nucleotide sites were obtained, including 111 in internal protein-coding genes which were significantly associated with mammalian virulence of multiple subtypes of IAV, and 26 in surface protein-coding gene N1 which were significantly associated with mammalian virulence of viral strains of HXN1 (Table S4). Low correlation was observed between these nucleotide sites (data not shown). In internal protein-coding genes, most mammalian virulence-related nucleotide sites (55 sites) were in PB1, followed by PB2, PA, NP, MP and NS. Table 2 listed the top 20 nucleotide sites in internal protein-coding genes which had the highest correlation with mammalian virulence (more nucleotide sites were listed in Table S4).

**Table 2.**
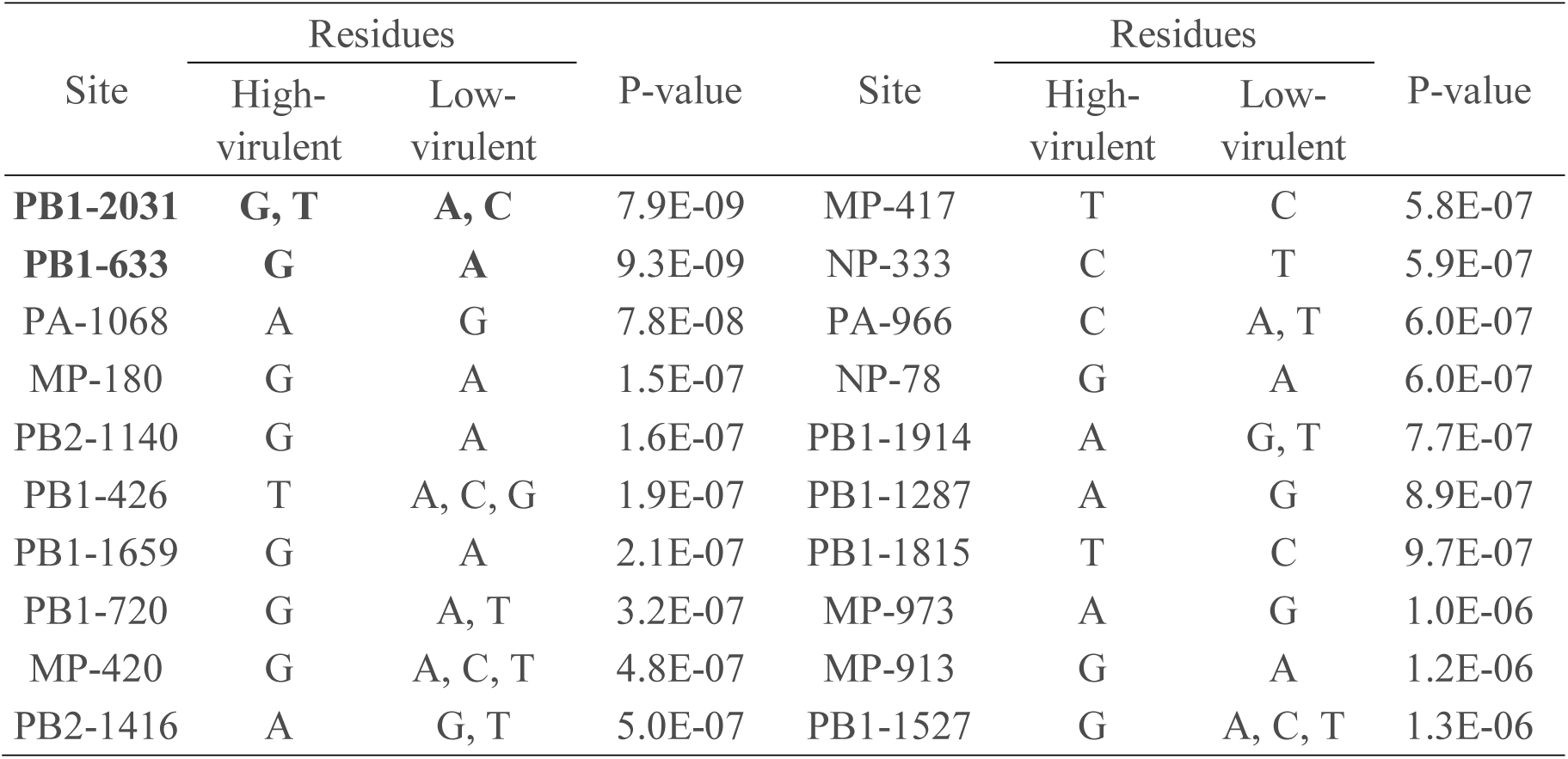
The top 20 nucleotide sites in internal protein-coding genes which were identified as mammalian virulence determinants. The top two sites for experimental validations were highlighted in bold. More details about these sites were provided in Table S4.

All of the 137 nucleotide sites were located in coding regions and 122 of them were located in the third position of codons. Further analysis showed that for 50 of these sites, the mutations between the low-virulent and high-virulent residues didn’t cause any amino acid replacement (Table S4), suggesting that these sites influenced the mammalian virulence totally at nucleotide level. For example, the top one nucleotide site (PB1-2031), had four kinds of nucleotides, all of which belonged to the codons encoding Threonine. For the remaining 87 nucleotide sites, their effects on mammalian virulence were mainly mediated at nucleotide level since the amino acid sites they encoded had no significant association with the virulence.

### 3.3 Validate the effects of synonymous mutations on mammalian virulence by *in vivo* experiments

To validate the nucleotide sites identified by association analysis, we investigated the influence of synonymous mutations (without changing amino acid) in the top two nucleotide sites, i.e., PB1-2031 and PB1-633, on the pathogenicity of IAV in mammals. According to the results above (Table 2), mutant viruses (denoted as m2031 and m633 respectively) with the mutation of A to G on both nucleotide sites were supposed to have higher virulence than the wild-type virus in mammals. They were used to infect the C57BL/6J mice with a dose of 10^6^ TCID_50_/50μl (see Methods). Body weight of each mouse was recorded daily, as well as PBS group mice. As shown in Fig. 2A, the wild-type pandemic H1N1 virus A/California/04/2009 (CA) caused the maximum reduction of body weight with 30% at 7 days post inoculation (dpi) in mice. Compared to the wild-type virus, the mice infected with the mutants (m2031, in red; m633, in blue) exhibited more obvious body weight loss (p-value < 0.01 in the paired t-test), although the trend of body weight loss was similar. Fig. 2B shows that mice infected with mutants died much earlier than those infected with wild-type viruses (3 or 4 dpi *vs* 6 dpi). In addition, 4/5 of the mice infected with the mutant m633 died as of 5 dpi, while 3/5 and 2/5 of the mice infected with the mutant m2031 and wild-type viruses died as of 7 dpi, respectively (Fig. 2B). For robustness of the results, we also tested the influence of these mutations in another H1N1 virus (A/Hunan/42443/2015, a human isolated Eurasian avian like swine influenza virus) with different genetic background. Similar results were obtained as those in genetic background of the CA virus (Fig. S3). Overall, these results verified that the synonymous mutations in PB1-2031 and PB1-633 could increase the pathogenicity of H1N1 viruses in mice.

**Fig 2.**
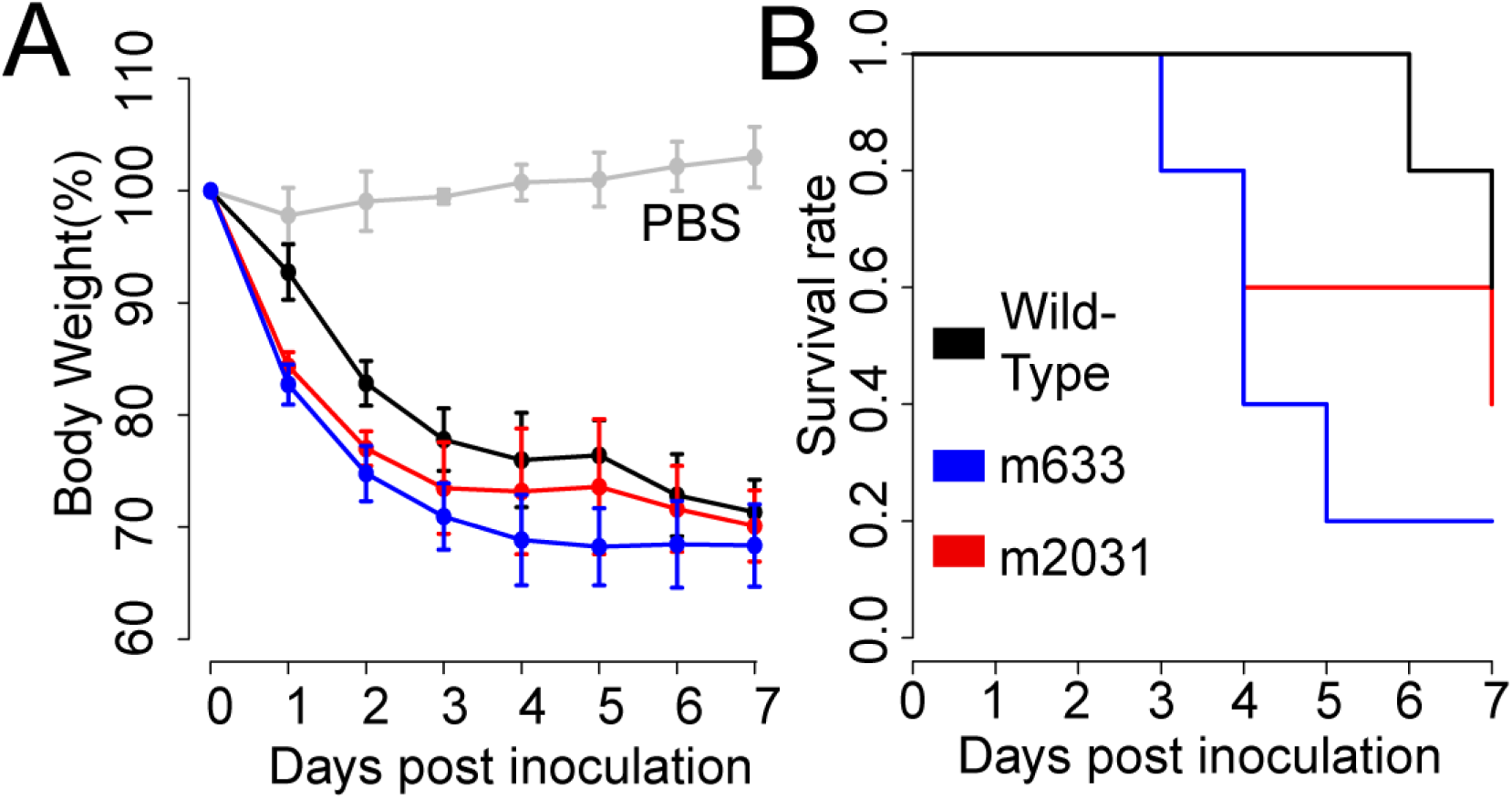
Experimental validation of the influence of the synonymous mutations in the top two nucleotide sites (PB1-2031 and PB1-633) on the *in vivo* features of IAV. (A) & (B) indicated body weight loss and mortality caused by the wild-type pandemic H1N1 virus A/California/04/2009 (CA) and the mutants in C57BL/6J mice, respectively. Wild-type viruses were colored in black, and the mutants m2031 and m633 were colored in red and blue, respectively. The error bar represented the standard deviation of five measurements.

### 3.4 Predicting mammalian virulence of IAV

Based on the nucleotide sites identified to be associated with mammalian virulence, we attempted to predict the mammalian virulence of IAV. Because of the difficulties in aligning genomic sequences of multiple subtypes of HA or NA, only 111 nucleotide sites in internal protein-coding genes were used. We firstly assessed the predictive ability of the single nucleotide site. PB1-720 was found to have the best predictive ability in discriminating between low-virulent and high-virulent strains. The model based on the site achieved an area under a receiver operating characteristic curve (AUC) of 0.65 and captured 35% of high-virulent strains (with sensitivity of 0.35) (Table 3). Then, we tried to combine multiple nucleotide sites in modeling. The naive bayes model had the best performance among multiple kinds of models in cross-validation test (Table S5). It achieved an AUC of 0.83 and captured 75% of high-virulent strains (Table 3).

**Table 3.** Performance of the models for predicting the mammalian virulence of IAV based on virulence determinants at the level of amino acid and nucleotide sites.

| Features | Models | Accuracy | Sensitivity | Specificity | AUC |
| --- | --- | --- | --- | --- | --- |
| Nucleotide sites | Single feature<br>(PB1-720) | 0.77 | 0.35 | 0.94 | 0.65 |
|  | Naive Bayes | 0.76 | 0.75 | 0.77 | 0.83 |
| Amino acid sites | Single feature<br>(PB2-627) | 0.75 | 0.4 | 0.90 | 0.65 |
|  | Naive Bayes | 0.73 | 0.75 | 0.72 | 0.84 |
| Combining nucleotide<br>and amino acid sites | Naive Bayes | 0.80 | 0.79 | 0.80 | 0.85 |

For comparison, we also assessed the ability of amino acid sites identified to have association with mammalian virulence in discriminating between high-virulent and low-virulent strains as above. PB2-627, the widely recognized molecular marker for virulence, was found to perform best among all 40 amino acid sites in internal proteins. The model based on it had a similar AUC with that of the nucleotide site PB1-720 (Table 3). Combination of multiple amino acid sites for modeling was conducted as above. The naive bayes model performed similarly with that based on nucleotide sites (Table 3 and Table S6).

Finally, to improve the predictive performances of models, we attempted to incorporate both amino acid and nucleotide sites identified above in modeling. The naive bayes method performed best compared to other methods (Table 3 and Table S7), with the values of AUC, predictive accuracy, sensitivity and specificity equal to 0.85, 0.80, 0.79 and 0.80, respectively. This model improved much in both the overall performance and sensitivity compared to those based on either amino acid or nucleotide sites (Table 3).

## 3 Discussion

In this study, we obtained genome-wide nucleotide sites associated with mammalian virulence by collecting hundreds of viral strains with diverse genetic background and mammalian virulence, and by computational analysis. More than 1/3 of these nucleotide sites were observed to only experience synonymous mutations. *In vivo* experiments showed that the synonymous mutations in either of the top two nucleotide sites enhanced the pathogenicity of the viruses in mice. Finally, machine-learning models with accepted accuracy for predicting the mammalian virulence of IAV were built, which provided a method for rapidly determining the virulence of newly emerging IAV.

Molecular markers play important roles in surveillance of newly emerging influenza viruses, such as monitoring the antigenic variation (Koel, et al., 2013), drug resistance (McKimm-Breschkin, 2013) and host adaptation of the virus (Chen, et al., 2006). Virulence is an important phenotype of the influenza viruses. Previous studies have identified lots of mammalian virulence-related molecular markers of the virus, such as PB2-627, PB1-317, NS1-92, and so on. However, all of them were identified at the amino acid level. Actually, mutations at the nucleotide level may also influence the pathogenicity of IAV. They can affect the packaging, transcription and translation of the virus, and interfere with the hosts’ immune response (Ferhadian, et al., 2018; Ingle, et al., 2015; Wang, et al., 2017; Williams, et al., 2018; Zhao, et al., 2014). This study was the first systematic analysis of the mammalian virulence determinants at nucleotide level for IAV. The nucleotide sites identified covered all internal protein-coding genes of IAV, which further validated the polygenic attribute of virulence. Interestingly, the PB1 gene, which is the RNA-dependent RNA polymerase (RdRP) subunit and responsible for synthesis of viral mRNA and genomic RNA, was found to have half of all nucleotide sites identified as virulence determinants. These nucleotide sites in PB1 may play important roles in transcription and translation of the PB1 protein. In addition, the nucleotide site in PB1 gene, i.e., PB1-720, was observed to have a similar ability to the widely used molecular marker PB2-627 in discriminating between low-virulent and high-virulent strains, suggesting its potential as a molecular marker of mammalian virulence for IAV.

Due to codon redundancy, mutations in the third position of codons are mostly synonymous. Most nucleotide-level virulence determinants identified here were located in the third position of codons. Over 1/3 of the nucleotide-level virulence determinants only experienced synonymous mutations. We firstly demonstrated that synonymous mutations in either of two nucleotide sites (PB1-2031 and PB1-633) could significantly influence the pathogenicity of IAV in mice, but the detailed mechanism was unknown.

Current molecular markers of viral virulence were mostly derived from small-scale experiments. They may work well in some viral strains or some subtype of IAV. Due to the widespread epistasis among amino acid sites, they may not work in the viral strains with different genetic background. For example, the mutation D701N in PB2 protein, which could enhance the pathogenicity of H9N2 virus in mice (Sediri, et al., 2016), had little impact on the pathogenesis of the 2009 pandemic H1N1 virus in both mice and ferrets (Jagger, et al., 2010). Computational identification of the molecular marker by integrating data from multiple sources or diverse genetic background may be benefit for recognizing those with more general impact. This is best exemplified in the GWAS studies, during which genetic variants identified with data from diverse population may be more useful for disease treatment (Frayling, et al., 2007; Ge, et al., 2009). The nucleotide-level virulence determinants identified here were obtained by analysis of more than 200 viral strains of 21 subtypes of IAV derived from nearly 100 studies. They may also work in IAV with diverse genetic background. For example, the mutations in PB1-633 and PB1-2031 had similar effects on the mammalian virulence in two viral strains of different genetic background (CA and HuN, Fig. 2 & Fig. S3). More researches are needed to confirm their influence on other subtypes of IAV. Anyway, the virulence determinants identified here can be helpful for prevention and control of the newly emerging IAV which are very probably to have different genetic background from the current viruses.

There were a few limitations in this study. Firstly, due to the research bias towards IAV widespread in the globe, the viral strains used in the present analysis were biased towards the subtypes of H5N1, H9N2, and so on. Bootstrap method was used to reduce the influence of this bias. The nucleotide sites with high bootstrap values may be more robust to the subtype of IAV; secondly, only two nucleotide sites identified as potential virulence determinants of IAV in this study were experimentally validated. The mechanism of nucleotide sites affecting viral pathogenicity was still not clear yet. More experiments were needed to validate the remaining candidate virulence determinants and clarify the mechanisms. Finally, the performance of computational models for predicting the mammalian virulence of IAV needs further improvement and integrating the contributions of HA and NA may help improve these models.

In summary, this work identified more than 100 mammalian virulence-related nucleotide sites in IAV. Over 1/3 of them experienced only synonymous mutations, highlighting the importance of synonymous mutations in mammalian virulence. Besides, the bio-markers identified and the computational models developed in the present study provided rapid and relatively accurate methods for evaluating the virulence of the virus, thus helping for early warning of newly emerging IAV.

